# Targeting PI3K-gamma in myeloid driven tumour immune suppression: A Systematic Review and Meta-Analysis of the Preclinical Literature

**DOI:** 10.1101/2024.05.12.593156

**Authors:** H Xu, SN Russell, K Steiner, E O’Neill, KI Jones

**Author notes:** Corresponding author: Keaton Ian Jones.

## Abstract

The intricate interplay between immune and stromal cells within the tumour microenvironment (TME) significantly influences tumour progression. Myeloid cells, including tumour-associated macrophages (TAMs), neutrophils (TANs), and myeloid-derived suppressor cells (MDSCs), contribute to immune suppression in the TME ^1,2^. This poses a significant challenge for novel immunotherapeutics that rely on host immunity to exert their effect. This systematic review explores the preclinical evidence surrounding the inhibition of phosphoinositide 3-kinase gamma (PI3Kγ) as a strategy to reverse myeloid-driven immune suppression in solid tumours.

EMBASE, MEDLINE, and PubMed databases were searched on 6^th^ October 2022 using keyword and subject heading terms to capture relevant studies. The studies, focusing on PI3Kγ inhibition in animal models, were subjected to predefined inclusion and exclusion criteria. Extracted data included tumour growth kinetics, survival endpoints, and immunological responses which were meta-analysed. PRISMA and MOOSE guidelines were followed.

A total of 36 studies covering 73 animal models were included in the review and meta-analysis. Tumour models covered breast, colorectal, lung, skin, pancreas, brain, liver, prostate, head and neck, soft tissue, gastric, and oral cancer. The predominant PI3Kγ inhibitors were IPI-549 and TG100-115, demonstrating favourable specificity for the gamma isoform. Combination therapies, often involving chemotherapy, radiotherapy, immune checkpoint inhibitors, biological agents, or vaccines, were explored in 81% of studies. Analysis of tumour growth kinetics revealed a statistically significant though heterogeneous response to PI3Kγ monotherapy, whereas the tumour growth in combination treated groups were more consistently reduced. Survival analysis showed a pronounced increase in median overall survival with combination therapy.

This systematic review provides a comprehensive analysis of preclinical studies investigating PI3Kγ inhibition in myeloid-driven tumour immune suppression. The identified studies underscore the potential of PI3Kγ inhibition in reshaping the TME by modulating myeloid cell functions. The combination of PI3Kγ inhibition with other therapeutic modalities demonstrated enhanced antitumor effects, suggesting a synergistic approach to overcome immune suppression. These findings support the potential of PI3Kγ-targeted therapies, particularly in combination regimens, as a promising avenue for future clinical exploration in diverse solid tumour types.

**Graphical abstract:** 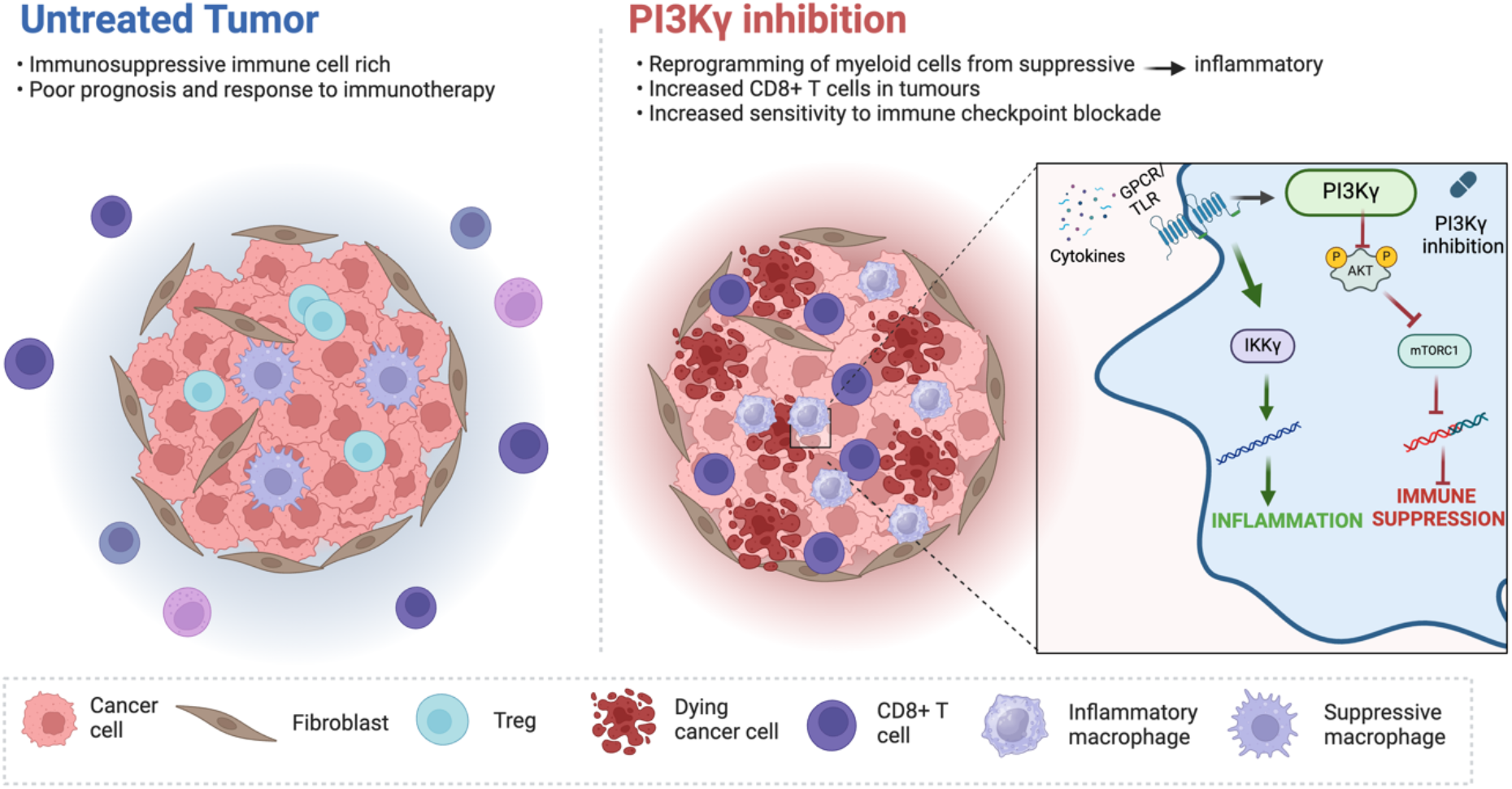

## Introduction

As our understanding of the complex tumour microenvironment (TME) has evolved, novel strategies that selectively target individual intertumoral components have been widely reported. The TME is comprised of immune and stromal cells, extracellular matrix, vascular networks, and cellular mediators alongside cancer cells. Host immune and stromal cells can be tissue resident or recruited from the systemic circulation and include macrophages, neutrophils, myeloid-derived suppressor cells (MDSCs), T lymphocytes, B lymphocytes, Natural Killer cells (NK), innate lymphoid cells (ILCs), and Fibroblasts. Cancer cells interact with these populations and can polarise their phenotypes in favour of tumour progression (i.e., tumour associated macrophage, TAM; tumour associated neutrophil, TAN; and cancer associated fibroblast, CAF) ^3–7^. Exploring the mechanisms of resistance to therapeutic strategies has underscored the critical contribution of the TME across a range of solid tumours. Foremost amongst these have been immune checkpoint inhibitors. Despite revolutionising the prognosis in select patient cohorts, checkpoint blockade remains ineffective in many solid tumour types ^8^. Both preclinical and clinical studies have demonstrated that myeloid cells (TAMs, TANs, MDSCs) appear to play a critical immune suppressive role within the TME ^9^. It is becoming clear that combinational approaches that target multiple components of the TME may be needed to overcome barriers to effective anti-tumour immunity. One of these approaches is the reversal of immune suppression by selectively targeting myeloid cells.

Decades of research has contributed to a comprehensive insight into the role of myeloid cells within the TME across multiple solid tumours. Myeloid populations comprising macrophages, monocytes (e.g., myeloid-derived suppressor cells), and granulocytes (e.g., neutrophils) have been shown to exert pro-tumourigenic functions. Expanding from the historic M1/M2 paradigm, tumoural macrophage phenotype has now been accepted as a continuum of heterogenous and pleotropic populations that respond to external selection pressure. This functional diversity means that unselective TAM depletion can deprive the TME of macrophages that are integral to promoting anti-tumour activity. Alternative strategies include those that aim to augment specific macrophage functions, such as phagocytosis^10^ and angiogenesis^11^, and those that reprogram macrophages through metabolic, epigenetic, and signalling pathways^12,13^. Reprogramming strategies are particularly attractive as they retain the potential benefits of inflammatory macrophages that may be an essential component of the anti-tumour immune cascade. One such approach is inhibition of the Phosphoinositide 3-Kinase gamma pathway.

Phosphoinositide 3-Kinase (PI3K) signalling plays a role in a wide range of biological processes. Class I PI3Ks are divided into class IA (comprising PI3K-α, β, and δ) and class 1B (consisting of PI3K-β). Isoforms PI3K-α and PI3K-β are expressed in a wide range of cells including epithelial cells, PI3K-δ is expressed in T lymphocytes, and PI3K-γ is uniquely expressed in myeloid cells ^14^. Activation of PI3K-γ leads to upregulation of signalling processes that are associated with an immune suppressive phenotype ^15^. Recent years have seen a large increase in the number of preclinical reports highlighting the role of PI3K-γ in tumour progression, as well as the effect of PI3K-γ inhibition using novel small molecule inhibitors. This can be attributed to the emergence of several isoform specific drugs that improved selectivity and avoided unwanted off target effects. In addition, there are now PI3K-γ inhibitors being tested in early phase clinical trials with promising results ^16^. We have systemically reviewed the current preclinical literature reporting on the effect of PI3K-γ inhibition in pre-clinical tumour models. Meta-analysis of tumour growth kinetic data, survival endpoints, as well as immunological responses provided valuable insights into the trends associated with this novel treatment strategy.

## Methods

### Search strategy

This review was conducted with reference to the Meta-analysis of Observational Studies in Epidemiology (MOOSE) and reported using the Preferred Reporting Items for Systematic Review and Meta-analyses (PRISMA). A qualified medical librarian conducted the literature searches on 6^th^ October 2022. The following databases were searched individually for relevant studies: Medline and Embase (both via Ovid) and PubMed (via National Library of Medicine). As the review was solely concerned with pre-clinical models, it was not deemed necessary to search trial registers. The search strategies included a combination of keyword searches and subject heading searches for synonyms of PI3Kγ and very broad subject heading and keyword terms for cancer. As there is a lot of inconsistency in how PI3Kγ is referred to and written (exacerbated by databases transliterating the Greek letter γ in inconsistent ways), many synonyms and a proximity search were necessary to create a sensitive search. Drug names, chemical designations, and commercial trademarked names were included. The full search strategies for each database can be viewed in the supplementary material. A total of 3128 results were retrieved from the searches (1970 after removal of duplicates). No search limits or filters were applied, for example study type, publication date or language. The results were deduplicated using EndNote, Deduklick (via risklick.ch), and Rayyan. Reference lists of key articles were hand-searched.

### Inclusion criteria

#### Population

Inclusion: Studies reporting the outcome of preclinical mouse or rat experiments.

Exclusion: Clinical trials and studies involving human subjects. Ex vivo studies. In vitro studies.

#### Intervention

Inclusion: The use of PI3Kγ selective inhibitors (can be delta/gamma isoform specific) as mono-or combinational therapy for cancer. All timings, dosages, frequencies, and administration routes.

Exclusion: Treatment involving Pan PI3K inhibitors (not described as gamma selective) or genetic depletion of PI3Kγ (e.g. knockout mice).

#### Comparator

Inclusion criteria: No treatment group, vehicle-treated group, or treatments that do no target PI3K in the case of combinational therapy Exclusion criteria: Healthy animals, animals that did not develop tumour where a tumour is expected

### Outcome

Inclusion criteria: Reported survival and/or tumour growth outcomes Exclusion criteria: No relevant outcomes reported

### Additional exclusion criteria

Include only publications in English. Exclude conference abstracts, posters, reviews, editorials, and theses.

### Procedure for study selection

Studies was selected through two rounds of screening. In the first round, only the title and abstract were screened. Selected articles then underwent full-text review in the second round of screening. Each round of screening was done by 2 reviewers independently, any discrepancies was resolved through discussion and consultation with the full study team of 4 reviewers.

### Data extraction methods

Data was tabulated by 2 reviewers and any discrepancies were discussed. Disease model, intervention method, dosage, schedule, survival outcome, and any immunological characterisations (by flow cytometry) were extracted for each model. For tumour growth data with fixed volumetric endpoint, time to reach end point was recorded, and vice versa. For median survival, time to reach 50% survival from Kaplan-Meier curves were recorded. GEM tumour models were excluded from survival data due to long incubation periods. Quantitative data was extracted from graphs through digital annotations.

### Data extraction

Extracted data included study design, treatment groups, control groups, disease model, species, method of cancer induction, intervention method, dosage, route, schedule, and survival outcome, survival data (i.e. disease free survival, progression free survival), tumour growth rate, intratumoural immune cell fraction (by flow cytometry), macrophage polarization status (assessed by classical M1/M2 markers), and bibliographical details.

### Statistical analysis

All statistical tests were performed using GraphPad Prism. Mixed-effects analysis (i.e. REML model with data matched within each publication) with Tukey’s multiple comparisons test was used for tumour volume, median survival, and stratified CD8 T cell data. Paired T test was used to compare CD8 T cell change between PI3Kγ inhibitor and Combo treatment. ns, not significant, ∗p ≤ 0.05, ∗∗p ≤ 0.01, ∗∗∗p ≤ 0.001, ∗∗∗∗p ≤ 0.0001.

## Results

### Study characteristics

Following removal of duplicates, a total of 1970 records were identified from searched databases, 36 of which were included in the review. Reasons for removal are summarised in the PRISMA flow diagram (Figure 1). The earliest publication across all included study was 2016. The tumour models used across these studies were breast (12), colorectal (12), lung (6), skin (6), pancreas (3), brain (2), liver (2), prostate (2), head and neck (1), soft tissue (1), gastric (1) and oral (1). Breast and colorectal models were used in 24/36 studies, with CT26 and 4T1 being the most commonly used cell lines. The most frequently used PI3K-gamma inhibitor was IPI-549 (Eganelisib, Infinity Pharmaceuticals). Mouse models were syngeneic in most cases. Tumour induction routes were subcutaneous (28/36), orthotopic (11/36), and genetically engineered (3/36). The most frequently used mouse strains were C57/BL6 (17) and BALB/c (17). 29 (81%) of the studies tested PI3K-γ inhibition with an additional agent. The combination therapy was chemotherapy (8), radiotherapy (3), immune checkpoint inhibitors (10), biological agents (3), and a vaccine (1).

**Figure 1.**
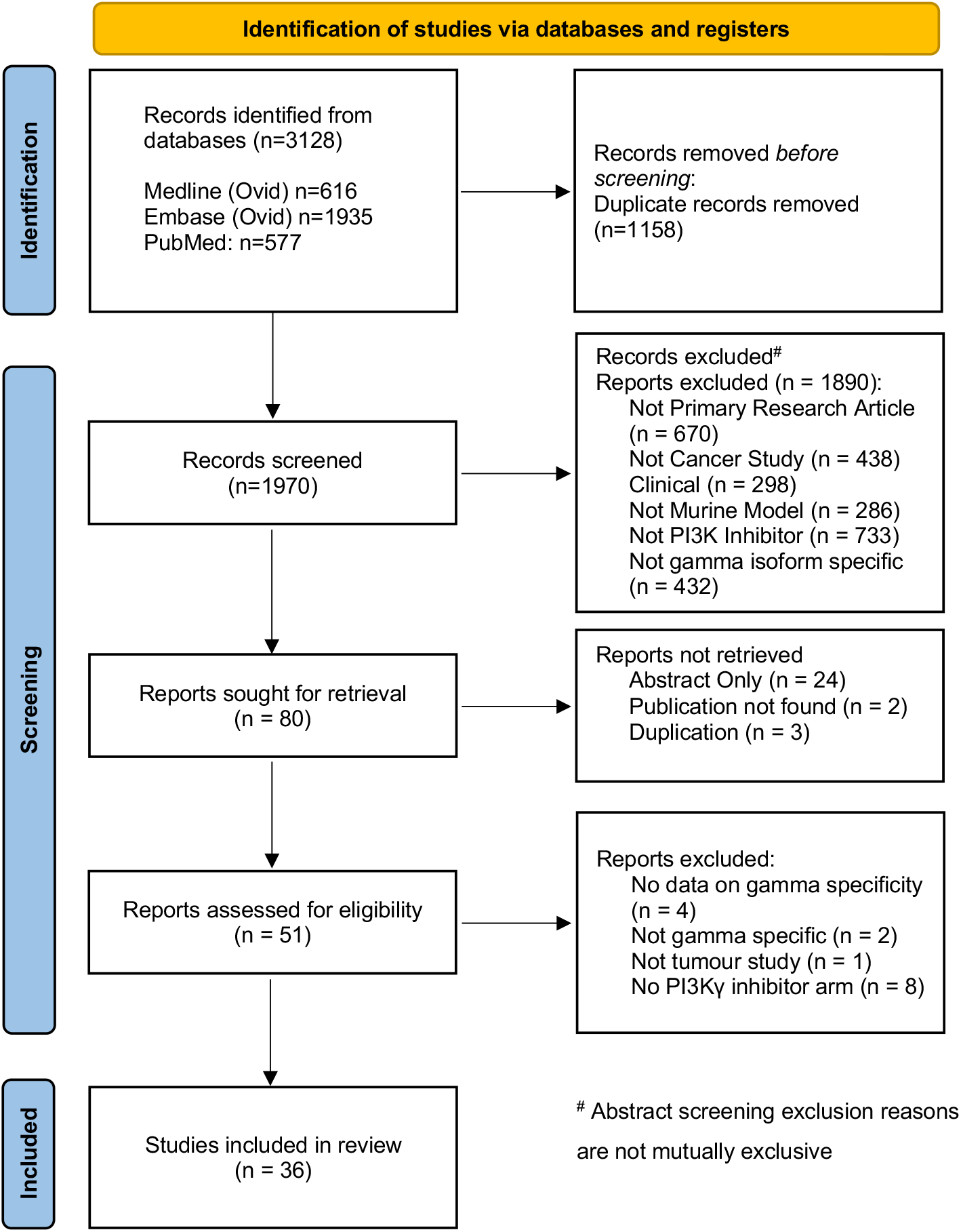
PRISMA flow diagram indicating the number of included studies and reasons for exclusion.

**Table 1.** Summary of the study characteristics of included publications.

|  | Reference | Inhibitor | Cancer Type | Cell Line | Model | Mouse Strain | Combo Type |
| --- | --- | --- | --- | --- | --- | --- | --- |
| 1 | (De Vera et al., 2019)<sup>17</sup> | IPI-549 | Colon | SW620 | SC | athymic NCR | Paclitaxel |
| 2 | (Zha et al., 2019)<sup>18</sup> | IPI-549 | Colon | CT26 | SC | Balb/c | CT-26 C3 KO |
| 3 | (Chung et al., 2020)<sup>19</sup> | AS-605240 | Prostate | PKC | GEM | PB-Cre4; p53; Kras |  |
| 4 | (Yoon et al., 2022)<sup>20</sup> | BR101801 | Colon | CT26 | SC | Balb/c | IR |
| 5 | (Qin et al., 2019)<sup>21</sup> | TG100-115 | Breast | 4T1 | SC | Balb/c | PLG-CA4 |
| 6 | (Yu et al., 2019)<sup>22</sup> | IPI-549 | Breast | 4T1 | SC | Balb/c | MnO2 nanoparticle |
| 7 | (Foubert et al., 2017)<sup>23</sup> | TG100-115 | Lung | LLC | SC | C57Bl/6J |  |
| 8 | (Li et al., 2018)<sup>24</sup> | IPI-549 | Breast | 4T1 | SC | Balb/c | doxorubicin |
| 9 | (Li et al., 2021)<sup>25</sup> | TG100-115 | Brain | CT-2A | ortho | C57BL/6 | temozolomide |
|  |  |  |  | GL261 | ortho | C57BL/6 |  |
| 10 | (Han et al., 2021)<sup>26</sup> | duvelisib (IPI-145) | Breast | 4T1-luc/PDX | SC | Balb/c/Huamanised | RT/PD-1 |
| 11 | (Martin et al., 2011)<sup>27</sup> | AS-605240 | Kaposi's Sarcoma | SV40 | SC | athymic mice | rapamycin |
| 12 | (Guan et al., 2022)<sup>28</sup> | IPI-549 | Colon | CT26 luc+ | SC | Balb/c | RT |
| 13 | (Wang et al., 2022)<sup>29</sup> | IPI-549 | Skin | B16F10 | SC | C57BL/6 | CpG (1mg/kg) in MOF Nanoparticle |
| 14 | (Ding et al., 2021)<sup>30</sup> | IPI-549 | Colon | CT26 | SC | Balb/c | photosensitizer |
| 15 | (Liu et al., 2022b)<sup>31</sup> | IPI-549 | Skin | B16F10 | SC | C57BL/6 | OVM |
|  |  |  | Colon | MC38 | SC | C57BL/6 | OVM |
|  |  |  | Pancreas | Pan02 | SC | C57BL/6 | OVM |
|  |  |  | Prostate | RM-1 | SC | C57BL/6 | OVM |
|  |  |  | Breast | 4T1 | SC | Balb/c | OVM |
| 16 | (Du et al., 2022)<sup>32</sup> | IPI-549 | Colon | MC38 | SC | C57BL/6J | Neoantigen vaccine |
| 17 | (De Henau et al., 2016)<sup>33</sup> | IPI-549 | Skin | B16-GMCSF | ID | C57BL/6J | anti-PD-1/CTLA-4 |
|  |  |  | Breast | 4T1 | SC | Balb/c | anti-PD-1/CTLA-4 |
| 18 | (Chang et al., 2020)<sup>34</sup> | AS-605240 | Breast | MDA-MB-231 | ortho | SCID | Paclitaxel (10mg/kg) |
| 19 | (Liu et al., 2022a)<sup>35</sup> | TG100-115 | Liver | Hepa1-6 | ortho, | C57BL/6J | anti-PD-1 |
| 20 | (Shen et al., 2021)<sup>36</sup> | IPI-549 | Colon | CT26 | SC | Balb/c | Oxaliplatin |
| 21 | (Jiang et al., 2020)<sup>37</sup> | IPI-549 | Breast | 4T1 | Ortho | Balb/c | silibinin (5 mg/ kg, IV) |
| 22 | (Schmid et al., 2011)<sup>38</sup> | TG100-115 | Lung | LLC | SC | C57BL/6 |  |
|  |  | AS-605240 | Lung | LLC | SC | C57BL/6 |  |
| 23 | (Li and Zhao, 2019)<sup>39</sup> | TG100-115 | Liver | Hep-3B | SC | nude | sorafenib (20 mg/kg) |
| 24 | (Davis et al., 2017)<sup>40</sup> | IPI-145 | Oral | MOC1 | SC | C57BL/6 | anti-PD-L1 |
| 25 | (Song et al., 2022)<sup>41</sup> | IPI-549 | Breast | MMTV-PyMT/4T1 | GEM/Ortho | MMTV-PyMT/Balb/c | anti-PD-1, Paclitaxel |
| 26 | (Zhang et al., 2019)<sup>42</sup> | IPI-549 | Pancreas | KPC | Ortho | C57BL/6 |  |
|  |  |  | Skin | BPD6 | SC | C57BL/6 |  |
| 27 | (Miyazaki et al., 2020)<sup>43</sup> | IPI-549 | Brain | TMZ-resistant TS | SC | C57BL/6 | anti-PD-L1 |
| 28 | (Kaneda et al., 2016a)<sup>44</sup> | TG100-115 | Pancreas | KPC | Ortho/GEM | C57BL6/FVB | Gemcitabine, anti-PD-1 |
| 29 | (Luo et al., 2020)<sup>45</sup> | IPI-549 | Gastric | MFC | ID | 615 |  |
| 30 | (Carnevali et al., 2021)<sup>46</sup> | AZD3458 | Colon | MC38 | SC | C57/Bl6 | anti-CTLA-4/PD-L1/PD-1 |
|  |  |  | Colon | CT26 | SC | Balb/c | anti-PD-L1 |
|  |  |  | Breast | 4T1 | Ortho | Balb/c | anti-PD-1 |
| 31 | (Xu et al., 2022)<sup>47</sup> | IPI-549 | Lung | Stem cells | SC | Nude athymic BALB/c | EGFR inhibitor |
| 32 | (Li et al., 2022)<sup>48</sup> | IPI-549 | Colon | CT26 fLuc+ | ID, | Balb/c | anti-PD-L1 |
|  |  |  | Breast | 4T1 | SC | Balb/c | anti-PD-L1 |
| 33 | (Han et al., 2022)<sup>49</sup> | IPI-549 | Skin | B16/F10 | Forced Met, SC tumor, | C57/BL6J | GFE1 peptide targeting exsome |
| 34 | (Lee et al., 2020)<sup>50</sup> | TG100-115 | Colon | CT26 | SC | Balb/c |  |
| 35 | (Kaneda et al., 2016b)<sup>51</sup> | IPI-549 TG115-110 | Head and Neck | HPV+ HNSCC | SC | C57Bl/6J |  |
|  |  |  | Lung | LLC | SC | C57Bl/6J |  |
|  |  |  | Breast | PyMT | Ortho | C57Bl/6J |  |
|  |  |  | Skin | HPV- HPV+ | SC | C3He/J | Anti-PD1 |
| 36 | (Joshi et al., 2020)<sup>52</sup> | IPI-549 | Lung | LLC | SC | C57/BL6 |  |

### PI3Kγ inhibitors

A total of 6 different PI3Kγ inhibitors were used across the studies. The most frequently reported were IPI-549 (21/36) and TG100-115 (9/36). All of the drugs included in the studies reported favourable specificity for the gamma isoform (IC50 7.9nM – 83nM). IPI-549 is the only drug used to have been tested in humans with an acceptable safety profile, including in combination with anti-PD1 ^16^. IC50 values for the other PI3K isoforms were reported where applicable and tended to have higher activity against PI3K delta compared with alpha and beta. Notably, 11 studies reported the use of modifications aimed at enhancing delivery or to combine PI3Kγ inhibitors with other therapeutics ^22,28–30,36,37,39,41–43,48,49^. This most frequently involved the formation of nanoparticles loaded with additional tumour targeting agents. In some cases, these were conventional chemotherapeutic agents ^41^. Other examples include the use of Manganese Oxide based preparations which aim to alleviate tumour hypoxia ^22,28^, and photosensitising agents in combination with photodynamic therapy ^30^. IPI-549 was used in all reports with a nanoparticle delivery platform. Details of combination therapies including dose, route of administration and schedule are included in Supplementary Table 1.

### Tumour growth

Tumour volume reduced by an average of 37% in animals receiving PI3Kγ inhibition as a monotherapy, 48% in those receiving single agent comparator therapy, and 69% in those receiving combination treatment (Figure 2 A,B). Responses in the PI3Kγi monotherapy group were heterogenous with some studies reporting no effect compared to others demonstrating a profound effect on tumour growth. Interestingly, an absence of effect using PI3Kγ inhibition alone did not correlate with the effects seen with combined therapy. In studies reporting no reduction in tumour growth with PI3Kγi alone ^17,28,31,46^ a profound suppression was observed in the combination groups. Some studies reported minimal effect with both PI3Kγi and comparator monotherapy but a significant effect with combination treatment (22 Pan02 model, 11).

**Figure 2.**
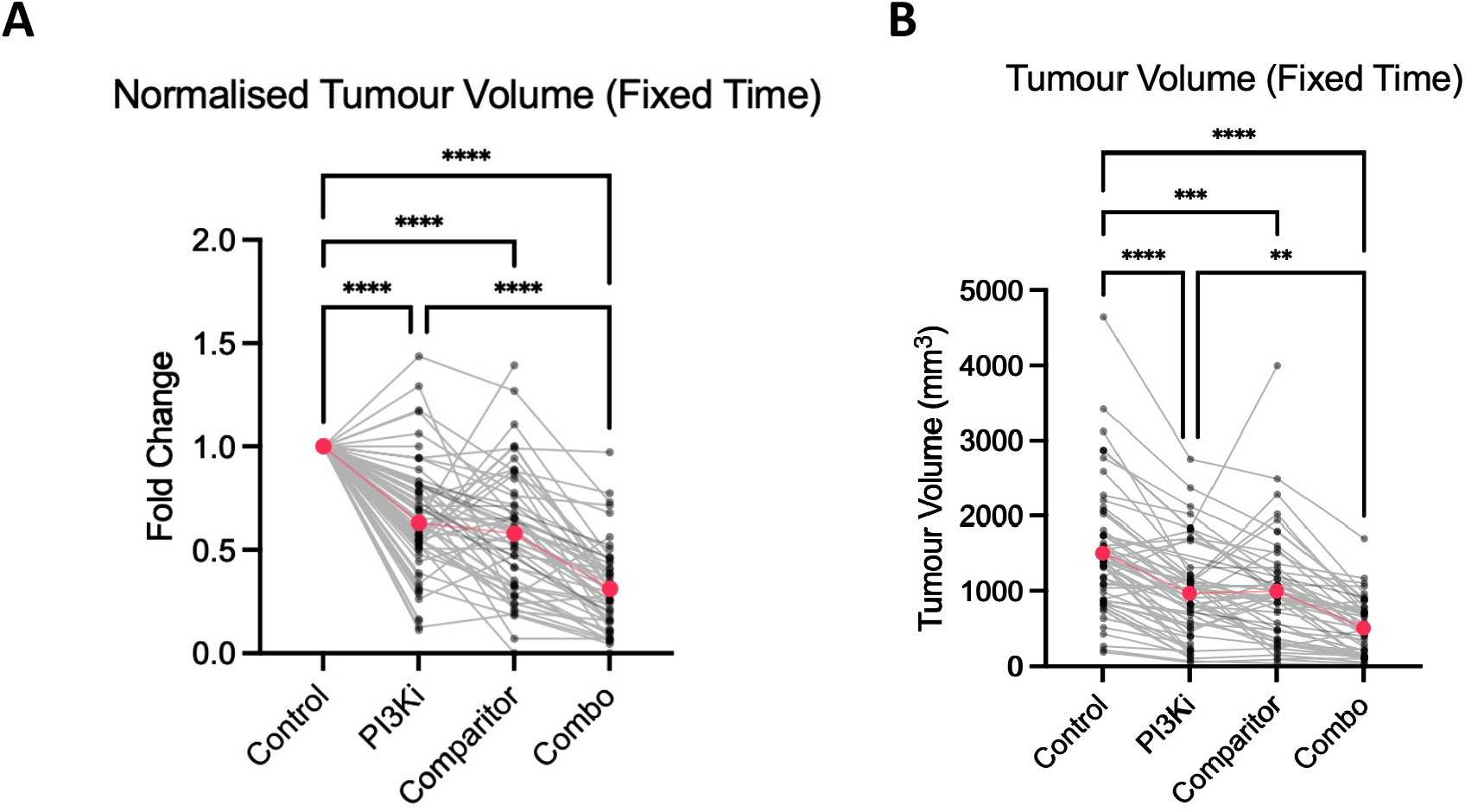
Changes in tumour growth in response to treatment. A, Tumour volume normalised to the average untreated tumour volume reported within each study. B, Absolute tumour volumes for each treatment group.

### Survival

Survival was reported in 25 studies. The median survival (OS, days) was 32.5, 35.5, 36.1 and 57.5 for control, PI3Kγ inhibition, comparator and combination cohorts respectively (Figure 3A). Compared to the differences observed in tumour growth kinetics, the effect of combination treatment on survival was more pronounced, with a relative increase in median OS of 15% and 26% in the PI3Kγ inhibitor and comparator groups compared with 81% in the combination group. In 4 studies median OS was not reached in the combination group ^22,26,28,53^, which was not observed in any of the other treatment groups across all studies. In these studies, the combination agent was an immune checkpoint inhibitor (anti-PD1/anti-PD-L1). Complete tumour regression (i.e., cure) was observed in 13 tumour models across 9 studies ^20,22,24–26,28,33,48,51^. Average rates of regression leading to long term survival were 0%, 1%, 6% and 26% in the untreated, PI3Kγ inhibitor, comparator and combination groups respectively (Figure 3B). It was noted that a minority of studies observed animals for recurrence for an extended period of time (>100 days, Figure 3C).

**Figure 3.**
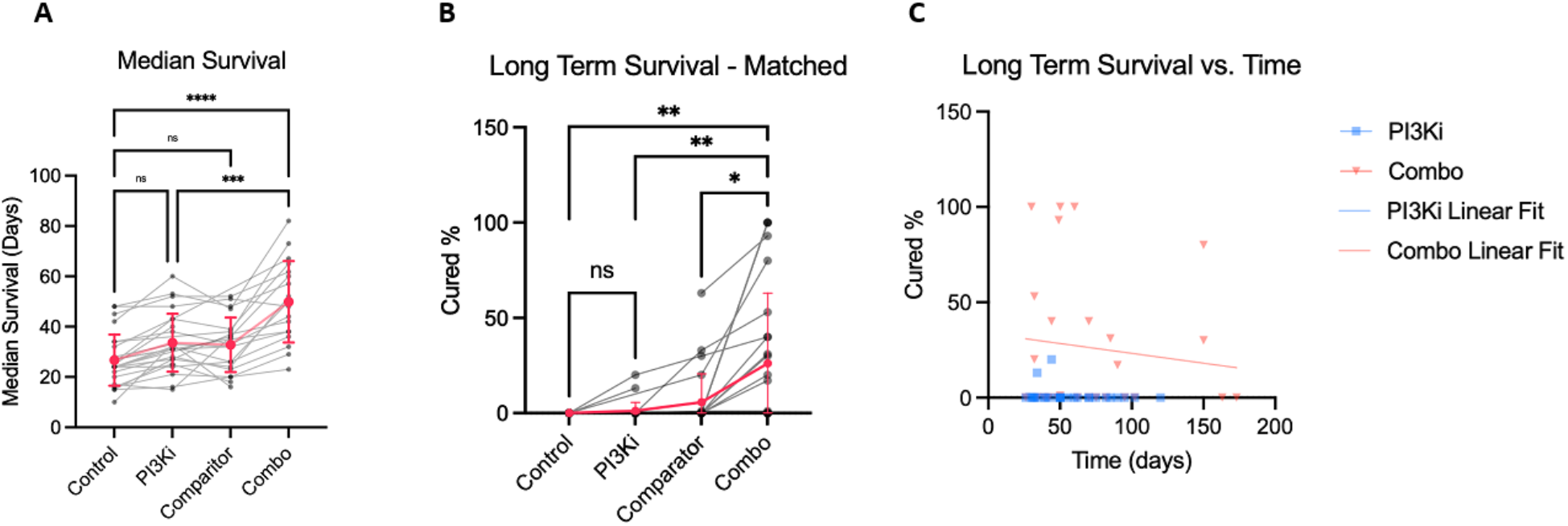
Summary of suvival data for groups receiving treatment as indicated. A, median oversall survival according to treatment group. B, rates of tumour regression (cure) according to treatment group. C, plot of cure rate over time with linear modelling according to PI3Kγ inhibitor monotherapy or combination treatment.

Additionally, 11/36 studies also reported metastatic burdens based on gross, histological, and radiological findings, and all of which demonstrated reductions in either PI3Kγi or combo treated groups ^19,21,22,29,36,41,44,48,49,51,53^. Breast and melanoma models accounted for over half of these reports (6/11) and included direct (i.v injection), orthotopic (4T1) and spontaneous (MMTV-PyMT) models. Two studies utilised dual tumour models (bilateral flank tumours) to demonstrate an abscopal effect with local treatment to a single tumour ^20,48^.

### Changes in immune cell populations

Most studies reported alterations in immune cell populations in tumours using techniques including flow cytometry, immunohistochemistry and RNA sequencing. The majority of studies focused on the myeloid compartment (macrophages, neutrophils and MDSCs) as well as lymphocytes (CD8, CD4). When comparing both PI3Kγ inhibitor and combination groups to controls, studies reported an increase in the proportion of M1 (inflammatory) macrophages and a decrease in M2 (suppressive) macrophages (Figure 4A,B). Other immune suppressive cells including MDSCs, neutrophils and regulatory T cells were also reduced in PI3Kγ treated groups. In 27 of the studies reporting on changes in CD8 T cells in the PI3Kγ inhibitor treated cohort, significant increases were observed in 19 (70%, Figure 4), rising to 91% (20/22) in the combination treatment cohorts. The magnitude of CD8 T cell increase was most substantial in the combination treatment groups (Figure 5 A,B). The combination therapies in the 5 studies reporting the most significant increase in T cells were anti-PD1 (3), oncolytic virus, and radiotherapy. There was no correlation between the magnitude of CD8 T cell increase and the effect on tumour growth inhibition (Supplementary Figure 1). We gathered date on dose scheduling of immune checkpoint inhibitors. In all cases, checkpoint inhibitors were initiated concurrently with PI3Kγ inhibitors, with the total number of doses ranging between three and seven (Supplementary Table 2). In addition to quantitation of T lymphocytes, a number of studies interrogated their activation status with a focus on CD8+ T cells. In the majority of cases, this involved protein quantification (flow cytometry, ELISA) of cytotoxic markers including granzyme B, perforin, interferon-γ, CD107a and TNF-alpha. Two studies used interferon-γ ELISPOT assays to demonstrate antigen specificity of CD8+ T cells ^19,40^

**Figure 4.**
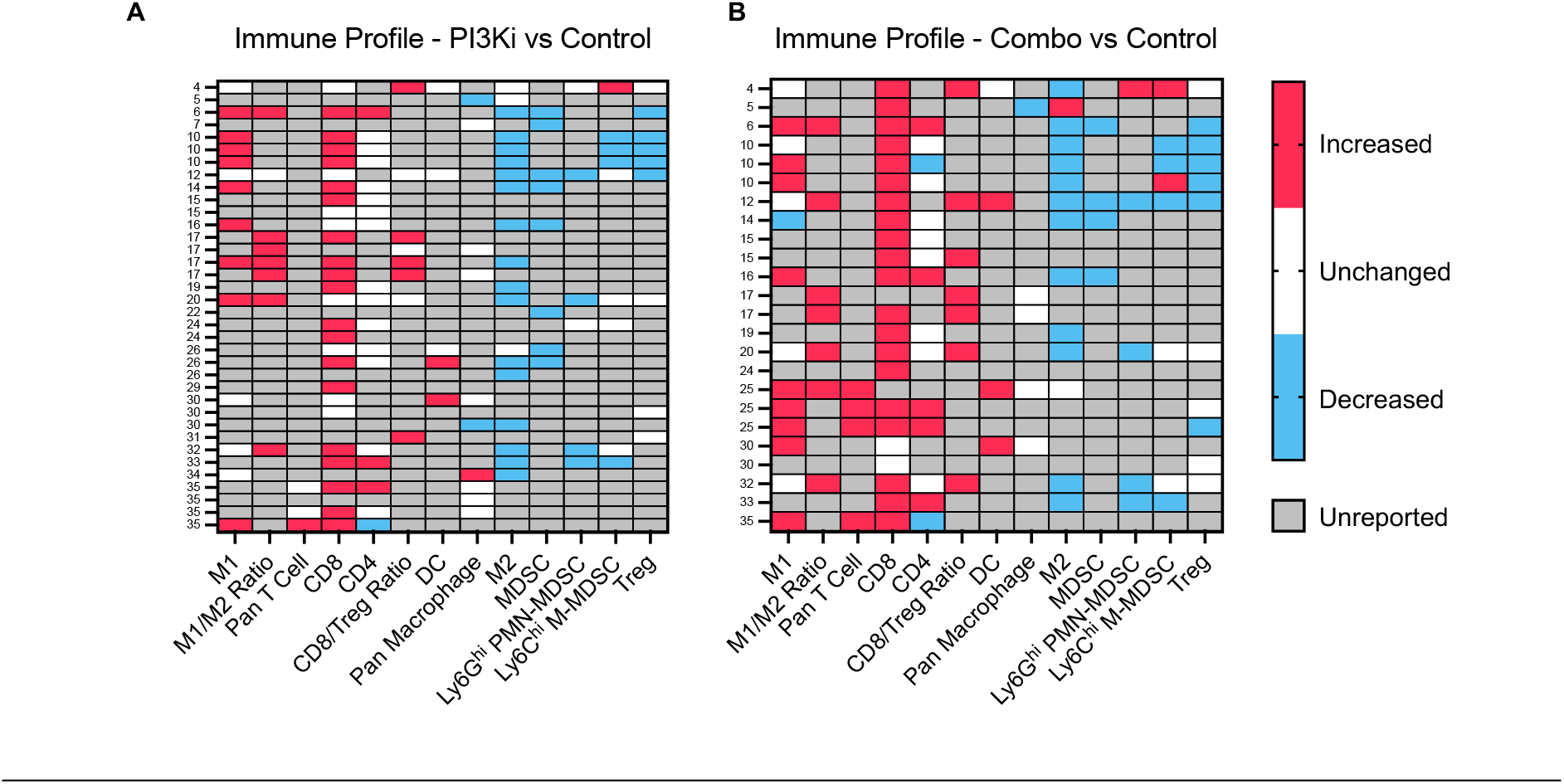
Heat maps showing relative change of immune cell populations reported by study. A, Immune fraction changes extracted from studies reporting the effect of PI3Kγ inhibition alone. B, Immune fraction changes extracted form studies comparing combination therapy to control. Row label refer to the study number as seen in Table 1.

**Figure 5.**
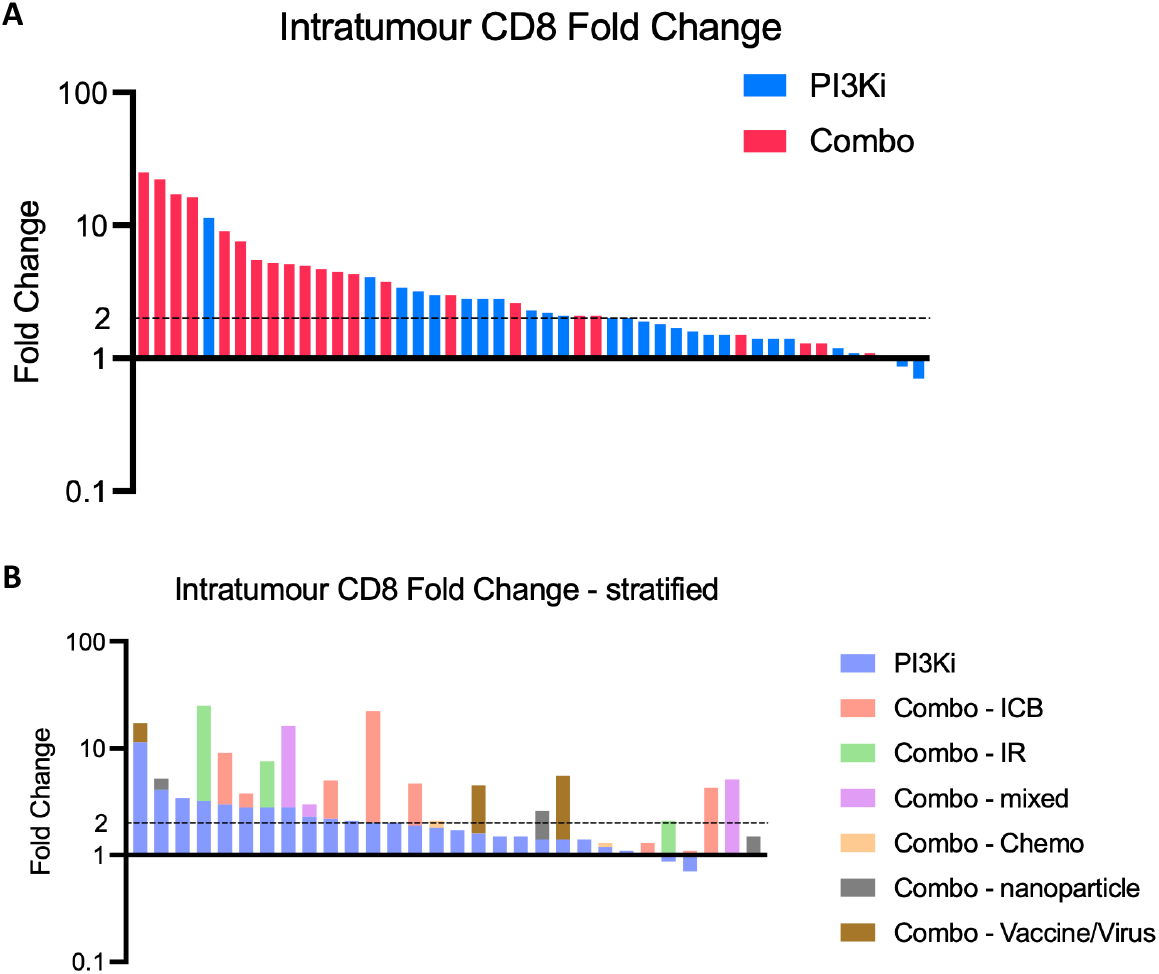
Changes in CD8 T cells according to treatment group. A, CD8 T cell fold change in groups receiving PI3Kγ inhibitor monotherapy (blue) and combination therapy (red). B, CD8 T cell fold change according to type of combination therapy.

Eight (8/36) of the studies described a non-immunological target for PI3Kγ inhibition^17,19,24,27,34,37,47,50^. This included direct targeting of tumour cells, increasing tumour cell sensitivity to chemotherapy, stem cells and stromal/extracellular matrix remodelling. One of these used an immune suppressed PDX model ^34^, the remaining were syngeneic. There was no significant difference in average changes in survival when immune versus non-immune studies were compared.

## Discussion

This systematic review of the preclinical literature summarises the current evidence supporting the use of PI3Kγ inhibitors for the treatment of solid tumours. Without specifying the inclusion of studies focusing on immune modulation a priori, we found that a minority (8/36) used PI3Kγ inhibitors with the aim of targeting tumour cells or other non-immune components. When tumour growth and survival data were meta-analysed, PI3Kγ when combined with any additional treatment had a more pronounced effect than monotherapies. This was particularly the case for combined treatment with immune checkpoint inhibitors. These effects were seen across tumour models and using different PI3Kγ inhibitors. Taken together, these consistent results demonstrate that PI3Kγ represents a promising target for clinical translation.

Combination treatment strategies have gained traction in light of the heterogenous response to novel immunotherapeutics observed in clinical studies. Preclinical work has highlighted the contribution of macrophages to intra-tumoural immune suppression, limiting the effect of drugs relying on cytolytic T cell activity ^54–57^. Macrophage pleiotropy and the role that functional subsets may play in promoting anti-tumour immunity has shifted the focus away from depletion and towards reprogramming. This approach takes advantage of the anti-tumour TAM functions that may be essential in permitting immune mediated tumour killing.

The predominant PI3Kγ inhibitors described in included studies, notably IPI-549 and TG100-115, are reported to have favourable specificity for the gamma isoform ^58,59^. This, combined with the myeloid specificity of PI3Kγ, translates into a highly targeted therapy with minimal off target effects. The inclusion of various solid tumour models in reported studies, including breast, colorectal, lung, skin, pancreas, brain, liver, prostate, head and neck, soft tissue, gastric, and oral cancers, underscores the broad applicability of PI3Kγ-targeted strategies. This is in keeping with clinical data that suggests a pro-tumoural role for macrophages in most tumour settings ^60–64^.

Combination therapies emerged as a recurrent theme, featured in 81% of the studies. These combinations spanned chemotherapy, radiotherapy, immune checkpoint inhibitors, biological agents, and vaccines, reflecting a multifaceted approach to counter myeloid-driven immune suppression. We did not identify a single group of tumour models where PI3Kγ inhibition had a more pronounced effect compared with combination therapies.

Notably, the heterogeneous response to PI3Kγ monotherapy in tumour growth kinetics suggested the need for nuanced considerations in selecting optimal treatment regimens. However, the pronounced reduction observed in tumour growth with combination therapies highlights the potential synergistic effects when PI3Kγ inhibition is integrated into broader treatment strategies. It was clear that most studies rationalised the use PI3Kγ inhibitors due to their capacity to reverse myeloid driven immune suppression. This is likely to be the reason that the most frequently used combination treatment was immune checkpoint inhibitor therapy. The reduced tumour growth rate observed with combination treatment translated into improved survival in some studies. The relative increase in median overall survival was notably higher in the combination group compared to both PI3Kγ monotherapy and comparator groups. This trend suggests that the synergistic effects observed in tumour growth kinetics translate into significant improvements in survival outcomes. This was further emphasised by the finding that only groups receiving combination treatment achieved tumour regression and ‘cure’ across all studies.

The immunological landscape within the TME underwent notable transformations upon PI3Kγ inhibition. Studies focused on alterations in myeloid cell populations encompassing macrophages, MDSCs, dendritic cells and neutrophils, as well as regulatory T cells. Alterations in NK cell populations were rarely reported. The majority of studies reported both quantitative and functional changes in these populations using techniques including flow cytometry, RNA sequencing, immunohistochemistry, and suppression assays. For myeloid characterisation, changes in the absolute number of macrophages were variable, but more consistent were the reported increases in M1 macrophages and decreases in M2 macrophages. This was commonly based on the expression of CD80 and iNOS for M1, and CD206 and arginase for M2. Gene expression studies, particularly RNA sequencing, provided additional clarity on the potential reprogramming in response to PI3Kγ inhibition. Gene set enrichment analyses highlighted changes to inflammatory and suppressive pathways as expected, but also additional pathways including antigen presentation, metabolism and phagocytosis ^18,26,33,35,46^. In addition to macrophages, a number of studies specifically investigated the effect of treatment on MDSCs ^22,23,26,28,30,31^. Similar to the effects observed in macrophages, PI3Kγ inhibition reduced phosphorylation of AKT, a recognised downstream signal of PI3K activation. This corresponded with reduced expression of immune suppression markers and also T cell suppression. Amongst the studies that reported a decrease in the total number of MDSCs, one demonstrated increased MDSC apoptosis in response to PI3Kγ inhibition ^30^. Finally, some studies highlighted the effect of treatment on regulatory T cell populations ^20,26,30,40,41^. This was particularly the case in studies using inhibitors with activity against PI3Kδ. In these studies, changes in regulatory T cell populations were accompanied by the previously discussed changes in myeloid cells. In the absence of regulatory T cell depletion models, it is however difficult to determine the relative contribution of these suppressive immune cell populations within the TME.

Importantly, our analysis revealed a substantial increase in CD8 T cell populations, particularly in the combination treatment cohorts. These T cells were frequently reported as having high expression of effector markers including interferon-γ, TNA-α, granzyme B, and perforin. We observed no correlation between the magnitude of CD8 T cell increase and the changes in tumour volume, suggesting that the quality of T cells (i.e. antigen specific, activated) is critical for anti-tumour activity. To discern the mechanism behind increased T cell numbers, some studies highlighted the reduced proliferative suppression capacity of PI3Kγ inhibitor treated macrophages ^23,53,65^. These findings further support the rationale for combining PI3Kγ inhibitors with treatments that depend on adaptive anti-tumour immunity for therapeutic effect.

While most studies focused on immunological targets, a subset explored non-immunological aspects of PI3Kγ inhibition, such as direct targeting of tumour cells, increasing chemosensitivity, and alterations in the tumour microenvironment. De Vera *et al*. demonstrated that IPI-549 sensitised multi-drug resistant (P-gp overexpressing) cell lines to taxane based chemotherapy by inhibiting P-gp mediated drug efflux ^17^. The efficacy of combination therapy was more pronounced in P-gp overexpressed tumour. Chung *et al*. showed that PI3Kγ was expressed in a genetically engineered mouse model (GEMM) of prostate cancer (Trp53/KRAS^G12D^) where epithelial-to-mesenchymal transition was observed ^19^. Only 10% of tumour cells in this model expressed PI3Kγ and whilst inhibition *in vitro* has a profound effect, this did not translate in the *in vivo* setting. In another study, the authors reported the protective role for PI3Kγ inhibition in doxorubicin induced cardiotoxicity by augmenting cardiocyte autophagy ^24^. Treatment also led to a reversal of tumour immune suppression. In a rare subtype of breast cancer, one group reported a direct effect on tumour cells ^34^, and also indirectly by reducing TAM tumour cell EMT ^50^. These studies highlighted that PI3Kγ inhibition may have alternative functions outside of the immunological context in distinct tumour types.

Several studies utilized drug modifications, including nanoparticle delivery platforms, highlighting the innovation in drug delivery strategies to enhance the therapeutic efficacy of PI3Kγ inhibitors. The incorporation of these advanced delivery methods may address challenges related to drug bioavailability and distribution within the TME. They also permit dual targeting with additional agents that may ameliorate other barriers to anti-tumour immunity.

The robust preclinical evidence has driven the development of several clinical PI3Kγ inhibitors, with IPI-549 (Infinity Pharmaceuticals) being the most advanced to date. The results of a phase I/Ib trial combining IPI-549 with anti-PD1 were favourable and pave the way for phase 2 studies ^16^. The most significant translational challenge is likely to be the selection of candidate combination therapies as well as appropriate tumour settings.

In conclusion, the results of this systematic review underscore the potential of PI3Kγ inhibition as a promising approach to reverse myeloid-driven immune suppression in the TME. The synergistic effects observed in combination therapies, coupled with the modulation of immune cell populations, provide a compelling rationale for further clinical exploration. As the field advances, understanding the intricate interplay between PI3Kγ inhibition, immune modulation, and tumour-specific factors will be crucial for optimizing therapeutic strategies tailored to diverse solid tumour types. Future clinical trials driven by the findings of these preclinical studies hold the potential to unlock new dimensions in cancer immunotherapy.

## Supporting information

Search strategies

## Notes

### Competing Interest Statement

The authors have declared no competing interest.

