## Supplementary Table 1 for "Targeting PI3K-gamma in myeloid driven tumour immune suppression: A Systematic Review and Meta-Analysis of the Preclinical Literature"

| Reference | Inhibitor | *in vivo* Dose | Route | Dosing Regimen | Cancer Type | Cell Line | Model | Mouse Strain | Combo Type |
| --- | --- | --- | --- | --- | --- | --- | --- | --- | --- |
| (De Vera et al., 2019) | IPI-549 | 3 mg/kg | IP | once per 3 days, 4 doses | Colon | SW620 | SC | athymic NCR | Paclitaxel |
|  |  |  |  |  | Colon | SW620/Ad300, ABCB1-overexpressing | SC | athymic NCR | Paclitaxel |
| (Zha et al., 2019) | IPI-549 | 15 mg/kg | PO | daily, day 7-17 | Colon | CT26 | SC | Balb/c | CT-26 C3 KO |
| (Chung et al., 2020) | AS-605240 | 20 mg/kg | IP | daily, 5 days a week for 5 weeks | Prostate | PKC | GEM | PB-Cre4; p53; Kras |  |
| (Yoon et al., 2022) | BR101801 | 50mg/kg | PO | daily, 32 days | Colon | CT26 | SC | Balb/c | -        IR (2 Gy) |
|  |  |  |  |  |  |  |  |  | -        IR (7.5 Gy) |
| (Qin et al., 2019) | TG100-115 | 2.5 mg/kg | IP | twice daily | Breast | 4T1 | SC | Balb/c | PLG-CA4, Vascular disrupting agent |
| (Yu et al., 2019) | IPI-549 | 25 μg/mouse, Nanoparticle | IV | once per 2 days | Breast | 4T1 | SC | Balb/c | MnO2 nanoparticle |
| (Foubert et al., 2017) | TG100-115 | 2.5 mg/kg | IP | twice daily, for 14 days | Lung | LLC | SC | C57Bl/6J |  |
| (Li et al., 2018) | IPI-549 | 5 mg/kg | PO | daily, 3 weeks | Breast | 4T1 | SC | Balb/c | doxorubicin (4 mg/kg) |
|  | AS-605240 | 5 mg/kg | IP | daily, 3 weeks | Breast | rHER-2/NeuT | GEM | BALB/c rHER-2/NeuT | doxorubicin (4 mg/kg) |
| (Li et al., 2021) | TG100-115 | 2.5 mg/kg | IP | twice daily, 2-3 weeks | Brain | CT-2A | ortho | C57BL/6 | temozolomide (50 mg/kg) |
|  |  |  |  |  |  | GL261 | ortho | C57BL/6 |  |
| (Han et al., 2021) | duvelisib (IPI-145) | 15 mg/kg | IP | once per two days, for 2 weeks | Breast | 4T1-luc | SC | Balb/c | -        RT (24 Gy in 3 fractisons) |
|  |  |  |  |  |  |  |  |  | -        anti-PD-1 (10 mg/kg, IP, every 2 days for two weeks) |
|  |  |  |  |  |  |  |  |  | -        RT + anti-PD-1 |
|  |  |  |  |  |  | 4T1 | SC | Balb/c | -        RT (24 Gy given in 3 fractions, every 2 days for a week) |
|  |  |  |  |  |  |  |  |  | -        anti-PD-1 (10 mg/kg, IP, every 2 days for two weeks) |
|  |  |  |  |  |  |  |  |  | -        RT + anti-PD-1 |
|  |  |  |  |  |  | 4T1 | SC | Nude | -        RT (24 Gy given in three fractions, every 2 days for a week) |
|  |  |  |  |  |  |  |  |  | -        anti-PD-1 (10 mg/kg, IP, every 2 days for two weeks) |
|  |  |  |  |  |  |  |  |  | -        RT + anti-PD-1 |
|  |  |  |  |  |  | PDX | SC | Humanised HuNSG | -        RT (24 Gy given in three fractions, every 2 days for a week) |
|  |  |  |  |  |  |  |  |  | -        anti-PD-1 (10 mg/kg, IP, every 2 days for two weeks) |
|  |  |  |  |  |  |  |  |  | -        RT + anti-PD-1 |
| (Martin et al., 2011) | AS-605240 | 25 mg/kg | IP | daily | Kaposi's Sarcoma | SV40-immortalized murine endothelial cells expressing KSHV-vGPCR | SC | athymic mice | rapamycin (5 mg/kg/day i.p.) |
|  |  |  |  |  |  | SV40-immortalized murine endothelial cells expressing PyMT | SC | athymic mice | rapamycin (5 mg/kg/day i.p.) |
| (Guan et al., 2022) | IPI-549 | 1.5 mg/kg, Nanoparticle | IV | once per 2 days, 2 doses | Colon | CT26 luc+ | SC, post incomplete surgical resection | Balb/c | RT (2 successive 3 Gy at 8 h post-injection for two cycles) |
|  |  | 1.5 mg/kg, Nanoparticle + RT | IV | once per 2 days, 2 doses | Colon | CT26 luc+ | SC, Primary and Distal, post surgrical resection of primary | Balb/c | RT (2 successive 3 Gy at 8 h post-injection for two cycles) + aPD-L1 (3.75 mg/kg, ip, every two days, 2 doses) |
| (Wang et al., 2022) | IPI-549 | 5 mg/kg | IV | once per 2 days, 4 doses | Skin | B16F10 | SC | C57BL/6 | CpG (1mg/kg) in MOF Nanoparticle |
|  |  | 5 mg/kg, Nanoparticle | IV | once per 3 days, 4 doses | Skin | B16F10 | SC | C57BL/6 | CpG in Nanoparticle = 1 mg/kg, aPD-L1 ( 7.5 mg/kg IP) |
| (Ding et al., 2021) | IPI-549 | 3.0 mg/kg, liposome | iV | once per 2 days, 7 doses | Colon | CT26 | SC | Balb/c | photosensitizer chlorin e6 0.75 mg/kg |
| (Liu et al., 2022b) | IPI-549 | 300 mg/mouse | PO | daily, for 2 weeks | Skin | B16F10 | SC | C57BL/6 | OVM (1 x10^7 plaque-forming units (PFU)/mouse) i.v. daily 5 times |
|  |  |  |  |  | Colon | MC38 | SC | C57BL/6 | OVM (1 x10^7 plaque-forming units (PFU)/mouse) i.v. daily 5 times |
|  |  |  |  |  | Pancreas | Pan02 | SC | C57BL/6 | OVM (1 x10^7 plaque-forming units (PFU)/mouse) i.v. daily 5 times |
|  |  |  |  |  | Prostate | RM-1 | SC | C57BL/6 | OVM (1 x10^7 plaque-forming units (PFU)/mouse) i.v. daily 5 times |
|  |  |  |  |  | Breast | 4T1 | SC | Balb/c | OVM (1 x10^7 plaque-forming units (PFU)/mouse) i.v. daily 5 times |
| (Du et al., 2022) | IPI-549 | 15 mg/kg | PO | daily, for 2 weeks | Colon | MC38 | SC | C57BL/6J | peptidic microarchitecture trapped neoantigen vaccine with CpG (SC, 50 μg neoantigen peptide combined with 0.5 μg CpG, every 5 days, 4 doses) |
| (De Henau et al., 2016) | IPI-549 | 15 mg/kg | PO | daily, from day 7 to day 21 | Skin | B16-GMCSF | ID | C57BL/6J | -        anti-PD-1 (250 μg per mouse) |
|  |  |  |  |  |  |  |  |  | -        anti-CTLA-4 (100 μg per mouse) |
|  |  |  |  |  |  |  |  |  | -        anti-PD-1 (250 μg per mouse) & anti-CTLA-4 (100 μg per mouse) |
|  |  |  |  |  | Skin | B16 | ID | C57BL/6J |  |
|  |  |  |  |  | Breast | 4T1 | SC | Balb/c | -        anti-PD-1 (250 μg per mouse) |
|  |  |  |  |  |  |  |  |  | -        anti-CTLA-4 (100 μg per mouse) |
|  |  |  |  |  |  |  |  |  | -        anti-PD-1 (250 μg per mouse) & anti-CTLA-4 (100 μg per mouse) |
| (Chang et al., 2020) | AS-605240 | 18mg/kg | NA | NA | Breast | MDA-MB-231 | ortho | SCID | Paclitaxel (10mg/kg) |
| (Liu et al., 2022a) | TG100-115 | NA | NA | daily, for 10 days | Liver | Hepa1-6 | ortho, post insufficient radiofrequency ablation | C57BL/6J | anti-PD-1 |
| (Shen et al., 2021) | IPI-549 | 50 ug/mouse, hydrogel | SC | 1 dose | Colon | CT26 | SC | Balb/c | Oxaliplatin |
| (Jiang et al., 2020) | IPI-549 | 5 mg/kg | IV | once per 2 days, 7 doses | Breast | 4T1 | Ortho | Balb/c | silibinin (5 mg/ kg, IV) |
| (Schmid et al., 2011) | TG100-115 | 2.5mg/kg | IP | twice a day, for 21 days | Lung | LLC | SC | C57BL/6 |  |
|  |  | 0.25mg/kg | IP | twice a day, for 21 days | Lung | LLC | SC | C57BL/6 |  |
|  | AS-605240 | 2.5 mg/kg | IP | twice a day, for 21 days | Lung | LLC | SC | C57BL/6 |  |
| (Li and Zhao, 2019) | TG100-115 | 30mg/kg | IV | twice a week for 4 weeks | Liver | Hep-3B | SC | nude | sorafenib (20 mg/kg) |
| (Davis et al., 2017) | IPI-145 | 15 mg/kg | PO | daily, for 14 days | Oral | MOC1 | SC | C57BL/6 | anti-PD-L1 (200 mg/injection x3) |
|  |  | 50 mg/kg | PO | daily, for 14 days | Oral | MOC1 | SC | C57BL/6 | anti-PD-L1 (200 mg/injection x3) |
| (Song et al., 2022) | IPI-549 | 15 mg/kg | PO | daily | Breast with Lung mets | MMTV-PyMT | GEM | MMTV-PyMT transgenic (FVB/NJ) | anti-PD-1, 100 ug per mouse, 3 doses IP, day 66, 69, 72 |
|  |  | 5 mg/kg | PO | daily | Breast with Lung mets | MMTV-PyMT | GEM | MMTV-PyMT transgenic (FVB/NJ) | Paclitaxel (in albumin nanoparticle, 5 mg/kg, IV), anti-PD-1 (100 ug per mouse, IP) |
|  |  | 5 mg/kg | IP | once per 3 days, 5 doses | Breast with Lung mets | MMTV-PyMT | GEM | MMTV-PyMT transgenic (FVB/NJ) | Paclitaxel (in albumin nanoparticle, 10 mg/kg, IV), anti-PD-1 (100 ug per mouse, IP, 3 doses) |
|  |  | 5 mg/kg | PO | daily | Breast | 4T1 cells + RAW 264.7 cells (3:1 ratio) | Ortho | Balb/c | anti-PD-1 (100 ug/mouse, once per 3 days, 3 doses) |
|  |  | 5 mg/kg | IP | once per 3 days | Breast | 4T1 cells + RAW 264.7 cells (3:1 ratio) | Ortho | Balb/c | Paclitaxel (in albumin nanoparticle, 10 mg/kg, IV), anti-PD-1 (100 ug per mouse, IP, 3 doses) |
| (Zhang et al., 2019) | IPI-549 | 30 mg/kg | PO | NA | Pancreas | KPC | Ortho | C57BL/6 |  |
|  |  | 15mg/kg, Nanoparticle | IV | once per 3 days | Pancreas | KPC | Ortho | C57BL/6 |  |
|  |  | 15mg/kg, Nanoparticle | IV | once per 3 days | Skin | BPD6 | SC | C57BL/6 |  |
| (Miyazaki et al., 2020) | IPI-549 | 1 mg/kg | PO | daily, day 7 - day 16 | Brain | TMZ-resistant TS | SC | C57BL/6 | anti-PD-L1 (IP, 200 ng/mouse, every 3 days, 4 doses) |
| (Kaneda et al., 2016a) | TG100-115 | 2.5 mg/kg | IP | twice a day, from day 7 - day 21 | Pancreas | mCherry LMP | Ortho | C57BL6;129 |  |
|  |  |  |  |  |  | p53 2.1.1 | Ortho | FVB/n |  |
|  |  |  |  |  |  | p53 2.1.1 | Ortho | FVB/n | anti–PD-1 (100 μg/mouse, i.p., every 3 days, 5 doses) |
|  |  |  |  |  |  | LMP | Ortho | C57BL6;129 | gemcitabine (10 mg/kg, i.p., every four days, 6 doses) |
|  |  |  |  |  |  | KPC | GEM | KPC on C57/BL6 |  |
| (Luo et al., 2020) | IPI-549 | 15 mg/kg | PO | daily, from day 5 to day 15 | Gastric | MFC | ID | 615 |  |
| (Carnevalli et al., 2021) | AZD3458 | 20 mg/kg | PO | twice a day | Colon | MC38 | SC | C57/Bl6 | -        anti-CTLA-4 (10 mg/kg, 4 dose) |
|  |  |  |  |  |  |  |  |  | -        anti-PD-L1 (10 mg/kg, 4 dose) |
|  |  |  |  |  |  |  |  |  | -        anti-PD-1 (10 mg/kg, 4 dose) |
|  |  |  |  |  | Colon | CT26 | SC | Balb/c | anti-PD-L1 (10 mg/kg, 2 doses per week) |
|  |  |  |  |  | Breast | 4T1 | Ortho | Balb/c | anti-PD-1 (10 mg/kg, 2 doses per week) |
| (Xu et al., 2022) | IPI-549 | 5 mg/kg | NA | once per 2 days, for 12 days | Stem Cell - Lung Cancer Conditioned Media Polarised | mouse induced pluripotent stem cells polarized with LLC conditioned media | SC | Nude athymic BALB/c | EGFR inhibitor Gefitinib (100 mg/kg, once per two days) |
| (Li et al., 2022) | IPI-549 | 25 μg/mouse, hydrogel | peritumoral | once | Colon | CT26 fLuc+ | ID, post inadequate microwave ablation | Balb/c | anti-PD-L1 (50 μg per mouse, in hydrogel) |
|  |  |  |  |  | Breast | 4T1 | SC, post inadequate microwave ablation | Balb/c | anti-PD-L1 (50 μg per mouse, in hydrogel) |
| (Han et al., 2022) | IPI-549 | 0.2 mg/kg | IV | once per 2 days, 6 doses | Skin | B16/F10 | Forced Met, SC tumor, surgerical excision, then IV tumor injection | C57/BL6J | GFE1 peptide targeting exsome |
| (Lee et al., 2020) | TG100-115 | NA | IP | once per 2 days, from day 7 to day 21 | Colon | CT26 | SC | Balb/c |  |
| (Kaneda et al., 2016b) | IPI-549 | 15 mg/kg | PO | daily, from day 8 | Head and Neck | MEER HPV+ HNSCC | SC | C57Bl/6J |  |
|  |  |  |  |  | Lung | LLC | SC | C57Bl/6J |  |
|  |  |  |  |  | Breast | PyMT | Ortho | C57Bl/6J |  |
|  |  |  |  |  | Skin | SCCVII HPV- | SC | C3He/J |  |
|  | TG100-115 | 5 mg/kg | IP | twice per day, from day 1 | Skin | SCCVII HPV- | SC | C3He/J | anti-PD-1 (250 μg, every 3 days, from day 1) |
|  |  | 2.5 mg/kg | IP | twice per day, from day 6 | Skin | SCCVII HPV- | SC | C3He/J | anti-PD-1 (250 μg, every 3 days, from day 3) |
|  |  | 2.5 mg/kg | IP | twice per day, from day 11 | Head and Neck | MEER HPV+ HNSCC | SC | C57Bl/6J | anti-PD-1 (250 μg, 4 doses total, once every 3 days, from day 11) |
| (Joshi et al., 2020) | IPI-549 | 10 mg/kg | PO | five times a week, day 10 to day 21 | Lung | LLC | SC | C57/BL6 |  |
