## Supplementary Table 2 for "Targeting PI3K-gamma in myeloid driven tumour immune suppression: A Systematic Review and Meta-Analysis of the Preclinical Literature"

|  | Reference | Inhibitor | Cancer Type | Cell Line | Model | ICB Combination Type | ICB Combination Dosage | ICB Dose (Frequency) | ICB Route | PI3K vs ICB | CD8 Fold Change: PI3Ki vs Control | CD8 Fold Change: Combo vs Control |
| --- | --- | --- | --- | --- | --- | --- | --- | --- | --- | --- | --- | --- |
| 10 | (Han et al., 2021)12 | IPI-145 | Breast | 4T1-luc | SC | Anti-PD-1 | 10 mg/kg | 7 (every 2 days) | IP | Concurrent | 2.8 | 3.8 |
| 17 | (De Henau et al., 2016)19 | IPI-549 | Breast | 4T1 | SC | Anti-PD-1 | 250 µg/mouse | 4 (every 3 days) | IP | Concurrent, PI3Ki Continued for 5 more days | 3.0 | 9.1 |
| 19 | (Liu et al., 2022a)21 | TG100-115 | Liver | Hepa1-6 | Ortho | Anti-PD-1 | NA | 4 (every 3 days) | NA | Concurrent | 2.2 | 5.0 |
| 24 | (Davis et al., 2017)26 | IPI-145 | Oral | MOC1 | SC | Anti-PD-L1 | 200 µg/mouse | 3 (every 3 days) | IP | Concurrent, PI3Ki Continued for 9 more days | 1.9 | 4.7 |
| 25 | (Song et al., 2022)27 | IPI-549 | Breast | 4T1 | Ortho | Anti-PD-1 | 100 µg/mouse | 3 (every 3 days) | IP | Concurrent, PI3Ki Continued for >3 more days | NA | 4.3 |
| 30 | (Carnevalli et al., 2021)32 | AZD3458 | Colon  Colon | MC38 | SC | Anti-PD-1 | 10 mg/kg | 4 (NA) | IP | Concurrent, PI3Ki Continued for >4 more days | 0.7 | 1.1 |
|  |  |  |  | CT26 | SC | Anti-PD-L1 | 10 mg/kg | 4 (NA) | IP | Concurrent, PI3Ki Continued for >4 more days | 1.0 | 1.3 |
| 35 | (Kaneda et al., 2016b) 37 | TG115-110 | Skin | HPV- SCCVII | SC | Anti-PD-1 | 250 µg/mouse | 6 (every 3 days) | IP | Concurrent | 2.0 | 22.3 |
