## Supplementary Figure 1 for "Targeting PI3K-gamma in myeloid driven tumour immune suppression: A Systematic Review and Meta-Analysis of the Preclinical Literature"

### Slide 1
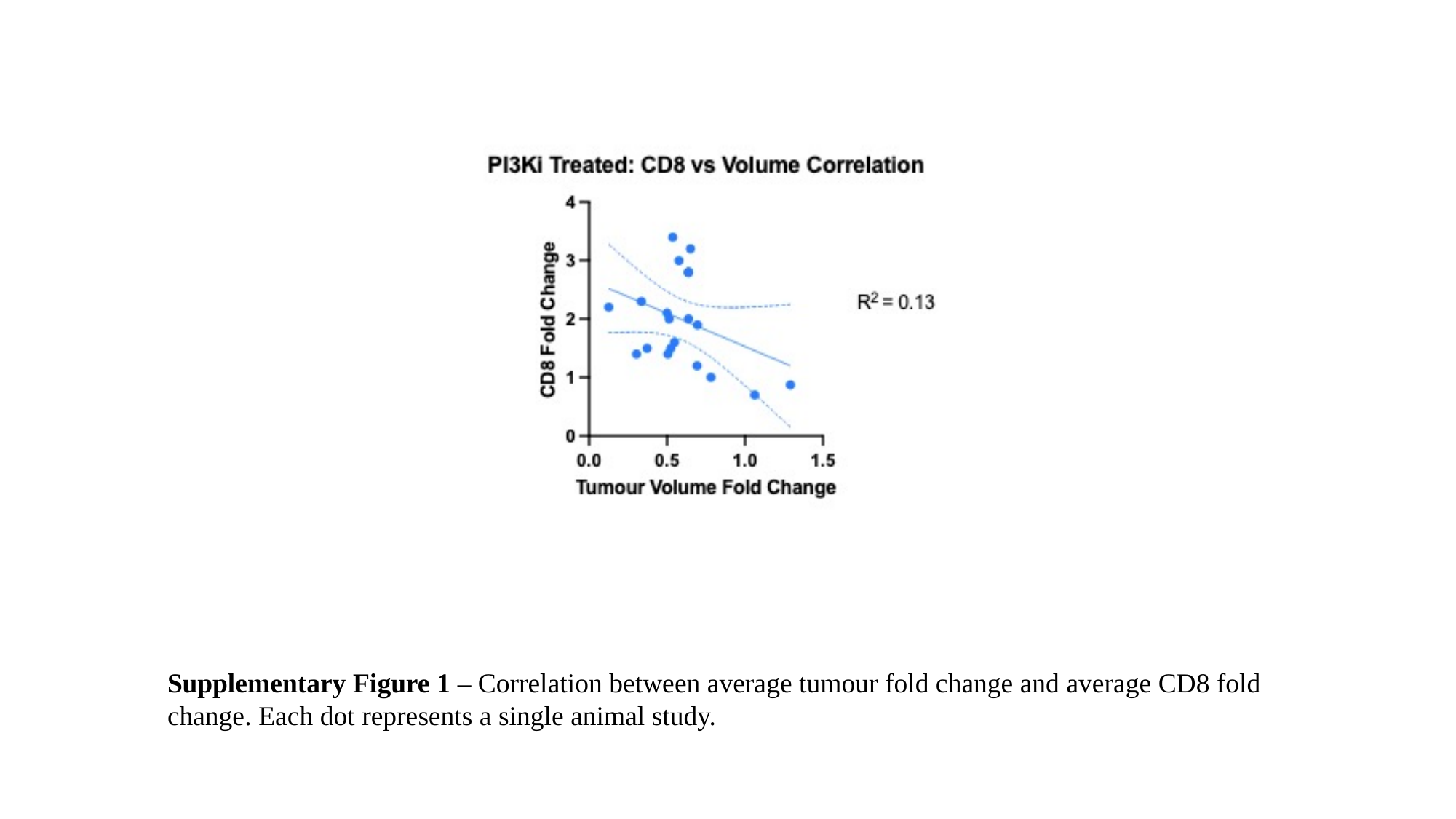

Supplementary Figure 1 – Correlation between average tumour fold change and average CD8 fold change. Each dot represents a single animal study.
