## Supplementary material for "Targeting PI3K-gamma in myeloid driven tumour immune suppression: A Systematic Review and Meta-Analysis of the Preclinical Literature": Search strategies

### Medline (Ovid MEDLINE® Epub Ahead of Print, In-Process & Other Non-Indexed Citations, Ovid MEDLINE® Daily and Ovid MEDLINE®) 1946 to present

06/10/2022

1	pi 3 k gamma.mp.

2	pi 3 kgamma.mp.

3	pi 3 kinase gamma.mp.

4	pi 3 kinasegamma.mp.

5	pi3 k gamma.mp.

6	pi3 kgamma.mp.

7	pi3 kinase gamma.mp.

8	pi 3k gamma.mp.

9	pi 3kgamma.mp.

10	pi3k gamma.mp.

11	pi3kgamma.mp.

12	pi3kinase gamma.mp.

13	pi3kinasegamma.mp.

14	phosphoinositide 3 kinase gamma.mp.

15	phosphoinositide 3 kinasegamma.mp.

16	phosphoinositide3 kinase gamma.mp.

17	phosphoinositide3 kinasegamma.mp.

18	phosphoinositide 3 oh kinase gamma.mp.

19	phosphatidylinositol 3 kinase gamma.mp.

20	phosphatidylinositol 3 kinasegamma.mp.

21	phosphatidylinositol 3 oh kinase gamma.mp.

22	phosphatidylinositol 3 oh kinasegamma.mp.

23	p13kgamma.mp.

24	RP6530.mp.

25	RP 6530.mp.

26	Tenalisib.mp.

27	Duvelisib.mp.

28	Copiktra.mp.

29	IPI145.mp.

30	IPI 145.mp.

31	IPI549.mp.

32	IPI 549.mp.

33	Eganelisib.mp.

34	ZX101A.mp.

35	ZX 101A.mp.

36	CAY10505.mp.

37	CAY 10505.mp.

38	AS605240.mp.

39	AS 605240.mp.

40	HM5023507.mp.

41	HM 5023507.mp.

42	BVD723.mp.

43	BVD 723.mp.

44	CAL130.mp.

45	CAL 130.mp.

46	TG100115.mp.

47	TG 100 115.mp.

48	AZD3458.mp.

49	AZD 3458.mp.

50	AS604850.mp.

51	AS 604850.mp.

52	(pi3k* adj4 gamma).mp.

53	PIK3CG.mp.

54	or/1-53

55	(cancer* or neoplasm* or tumour* or tumor*).mp.

56	exp Neoplasms/

57	55 or 56

58	54 and 57

### Embase 1974 to present

06/10/2022

1	pi 3 k gamma.mp.

2	pi 3 kgamma.mp.

3	pi 3 kinase gamma.mp.

4	pi 3 kinasegamma.mp.

5	pi3 k gamma.mp.

6	pi3 kgamma.mp.

7	pi3 kinase gamma.mp.

8	pi 3k gamma.mp.

9	pi 3kgamma.mp.

10	pi3k gamma.mp.

11	pi3kgamma.mp.

12	pi3kinase gamma.mp.

13	pi3kinasegamma.mp.

14	phosphoinositide 3 kinase gamma.mp.

15	phosphoinositide 3 kinasegamma.mp.

16	phosphoinositide3 kinase gamma.mp.

17	phosphoinositide3 kinasegamma.mp.

18	phosphoinositide 3 oh kinase gamma.mp.

19	phosphatidylinositol 3 kinase gamma.mp.

20	phosphatidylinositol 3 kinasegamma.mp.

21	phosphatidylinositol 3 oh kinase gamma.mp.

22	phosphatidylinositol 3 oh kinasegamma.mp.

23	p13kgamma.mp.

24	RP6530.mp.

25	RP 6530.mp.

26	Tenalisib.mp.

27	Duvelisib.mp.

28	Copiktra.mp.

29	IPI145.mp.

30	IPI 145.mp.

31	IPI549.mp.

32	IPI 549.mp.

33	Eganelisib.mp.

34	ZX101A.mp.

35	ZX 101A.mp.

36	CAY10505.mp.

37	CAY 10505.mp.

38	(pi3k* adj4 gamma).mp.

39	PIK3CG.mp.

40	AS605240.mp.

41	AS 605240.mp.

42	HM5023507.mp.

43	HM 5023507.mp.

44	BVD723.mp.

45	BVD 723.mp.

46	CAL130.mp.

47	CAL 130.mp.

48	TG100115.mp.

49	TG 100 115.mp.

50	AZD3458.mp.

51	AZD 3458.mp.

52	AS604850.mp.

53	AS 604850.mp.

54	1 or 2 or 3 or 4 or 5 or 6 or 7 or 8 or 9 or 10 or 11 or 12 or 13 or 14 or 15 or 16 or 17 or 18 or 19 or 20 or 21 or 22 or 23 or 24 or 25 or 26 or 27 or 28 or 29 or 30 or 31 or 32 or 33 or 34 or 35 or 36 or 37 or 38 or 39

55	or/1-53

56	(cancer* or neoplasm* or tumour* or tumor*).mp.

57	exp Neoplasm/

58	56 or 57

59	54 and 58

### PubMed

6/10/22

#1	"pi3kgamma"[All Fields] OR "pi 3 k gamma"[All Fields] OR "pi 3 kinase gamma"[All Fields] OR "phosphoinositide 3 kinase gamma"[All Fields] OR "p13kgamma"[All Fields] OR "phosphatidylinositol 3 kinase gamma"[All Fields] OR ("tenalisib"[Supplementary Concept] OR "tenalisib"[All Fields] OR "rp6530"[All Fields]) OR ("RP"[All Fields] AND "6530"[All Fields]) OR ("tenalisib"[Supplementary Concept] OR "tenalisib"[All Fields]) OR ("duvelisib"[Supplementary Concept] OR "duvelisib"[All Fields]) OR ("duvelisib"[Supplementary Concept] OR "duvelisib"[All Fields] OR "copiktra"[All Fields]) OR "IPI145"[All Fields] OR ("duvelisib"[Supplementary Concept] OR "duvelisib"[All Fields] OR "ipi 145"[All Fields]) OR "Eganelisib"[All Fields] OR "CAY10505"[All Fields] OR ("CAY"[All Fields] AND "10505"[All Fields]) OR ("ZX"[All Fields] AND "101A"[All Fields]) OR "PIK3CG"[All Fields] OR ("5 quinoxalin 6 ylmethylenethiazolidine 2 4 dione"[Supplementary Concept] OR "5 quinoxalin 6 ylmethylenethiazolidine 2 4 dione"[All Fields] OR "as605240"[All Fields]) OR "HM5023507"[All Fields] OR "TG100115"[All Fields] OR "AZD3458"[All Fields] OR ("5 2 2 difluorobenzo 1 3 dioxol 5 ylmethylene thiazolidine 2 4 dione"[Supplementary Concept] OR "5 2 2 difluorobenzo 1 3 dioxol 5 ylmethylene thiazolidine 2 4 dione"[All Fields] OR "as604850"[All Fields]) OR ("5 quinoxalin 6 ylmethylenethiazolidine 2 4 dione"[Supplementary Concept] OR "5 quinoxalin 6 ylmethylenethiazolidine 2 4 dione"[All Fields] OR "as 605240"[All Fields]) OR (("heart mind mumbai"[Journal] OR "hm"[All Fields]) AND "5023507"[All Fields]) OR ("BVD"[All Fields] AND "723"[All Fields]) OR ("cal 130"[Supplementary Concept] OR "cal 130"[All Fields] OR "cal 130"[All Fields]) OR (("trans gis"[Journal] OR "ieee trans games"[Journal] OR "tg"[All Fields]) AND "100"[All Fields] AND "115"[All Fields]) OR ("AZD"[All Fields] AND "3458"[All Fields]) OR ("5 2 2 difluorobenzo 1 3 dioxol 5 ylmethylene thiazolidine 2 4 dione"[Supplementary Concept] OR "5 2 2 difluorobenzo 1 3 dioxol 5 ylmethylene thiazolidine 2 4 dione"[All Fields] OR "as 604850"[All Fields])

#2	"neoplasms"[MeSH Terms] OR "cysts"[MeSH Terms] OR "cysts"[All Fields] OR "cyst"[All Fields] OR "neurofibroma"[MeSH Terms] OR "neurofibroma"[All Fields] OR "neurofibromas"[All Fields] OR "tumor s"[All Fields] OR "tumoral"[All Fields] OR "tumorous"[All Fields] OR "tumour"[All Fields] OR "neoplasms"[MeSH Terms] OR "neoplasms"[All Fields] OR "tumor"[All Fields] OR "tumour s"[All Fields] OR "tumoural"[All Fields] OR "tumourous"[All Fields] OR "tumours"[All Fields] OR "tumors"[All Fields] OR "cancer s"[All Fields] OR "cancerated"[All Fields] OR "canceration"[All Fields] OR "cancerization"[All Fields] OR "cancerized"[All Fields] OR "cancerous"[All Fields] OR "neoplasms"[MeSH Terms] OR "neoplasms"[All Fields] OR "cancer"[All Fields] OR "cancers"[All Fields]

#3	#1 AND #2
